# Soybean phenotypic plasticity in response to dark chilling stress

**DOI:** 10.1101/2024.05.07.592937

**Authors:** Chloé Elmerich, Michel-Pierre Faucon, Guénolé Boulch, Patrice Jeanson, Bastien Lange

## Abstract

The expansion of soybean cultivation to high-latitude regions is hindered by the adverse effects of dark chilling stress. This study investigates the phenotypic plasticity of soybean in response to dark chilling stress, aiming to identify traits that contribute to adaptation in northern climates. Five soybean cultivars from early maturity groups (000 and 00) were grown in growth chambers set either at 25/18°C or 25/10°C (day/night), and a range of performance and functional traits were measured. The relative distance plasticity index highlighted variations in plasticity across traits and genotypes. Photosynthetic activity recovery traits, shoot growth rate, and tap root elongation rate showed high plasticity and significant differences between cultivars, making them suitable for assessing dark chilling tolerance. Moreover, distinct responses to dark chilling stress were exhibited by different soybean cultivars. For instance, cv. SOPRANA displayed a greater phenotypic plasticity in aboveground performance traits, such as shoot growth, while cv. SULTANA exhibited a greater plasticity in physiological traits like photosynthetic activity recovery. These differing strategies may lead to similar biomass production through distinct mechanisms, highlighting the complexity of soybean’s adaptive responses to dark chilling stress. These findings provide valuable insights for breeders seeking to develop soybean cultivars adapted to high latitudes, meeting the increasing demand for locally produced protein-based food. However, further research in real field conditions is essential to validate the potential benefits of these plastic traits and their role in improving soybean resilience to dark chilling stress. The identified plastic traits offer a promising avenue for screening a broader range of cultivars and enhancing soybean’s suitability for cultivation in northern regions.

## 1. Introduction

Temperature, as an important evolutionary driver and a selective pressure factor in both natural selection and breeding, has consequences on the geographical distribution of species, including main crops like wheat, maize, rice, or soybean (Sloat et al., 2020). As low temperatures influence development rates, as well as delimiting the window of suitable conditions for plant growth, the potential to extend spring crops under high latitudes is limited (Menzel et al., 2006; Burrows et al., 2014). Although increases in temperature can be expected because of global warming, the selection of cultivars adapted to high latitudes (*i.e.,* ≥ 45°) is a major asset for this expansion northward (Maracchi et al., 2005; Elmerich et al., 2023a). At high latitudes, the vegetative growth of spring crops is especially affected by low minimum temperatures occurring at night (dark chilling). The ability of species or cultivars to tolerate low temperatures in the range of 0 to 15°C is referred to as chilling tolerance.

The increasing demand for locally produced protein-based food, for both animal and human consumption, has led to a growing interest in soybean (*Glycine max* [L.] Merill) cultivation in Europe (Boulch et al., 2021; Karges et al., 2022; Nendel et al., 2023). Similarly to many other spring crops such as maize or sorghum, soybean, a thermophilic short-day plant, is sensitive to chilling stress from germination to maturity (Cabané et al., 1992; Van Heerden et al., 2003; Ohnishi et al., 2010). Dark chilling is known to affect various plant physiological processes in different organs (*e.g*., leaves, roots, and seeds) and at different scales. Considering plant phenology, low temperatures (*i.e.*, below 15 °C) reduce the development rate, the emission of new leaves, and the duration of flowering and pod setting (Hume & Jackson, 1981). Additionally, low temperatures during reproductive growth increase flower abortion and pod abscission (Ohnishi et al., 2010). These traits presented different genotypic responses (Lawn & Hume, 1985; Seddigh et al., 1989). Moreover, photosynthetic activity is a crucial physiological process that is affected by low temperatures (Allen & Ort, 2001). Therefore, several parameters related to photosynthetic activity have been used to evaluate and compare the sensitivity of species and cultivars to low temperatures. These parameters include chlorophyll content (SPAD), chlorophyll fluorescence (F_v_/F_m_) and the quantum yield electron transport at the second photosystem (ΦPSII), which are considered as a sensitive indicator of resistance to a broad range of environmental stressors (Kautz et al., 2014). In soybean, Van Heerden & Krüger (2000) demonstrated severe mesophyll limitation of photosynthesis after a single night of low temperature exposure. Differences in genotypic response and sensibility to dark chilling were also observed when studying photosynthesis-related traits (Neuner & Larcher, 1990; Van Heerden et al., 2003). Finally, concerning belowground traits, although they have been reported in maize to be impacted by low temperatures, there is minimal understanding on the roots’ ecophysiological responses to dark chilling stress, their consequences on other physiological mechanisms (*e.g.*, nutrient and water uptake), and differences between cultivars (Hund et al., 2008). Dark chilling ultimately affects both functional traits and plant performance. Despite significant advances in understanding the dark chilling effects on soybean physiology, the relation between performance and functional traits including simultaneously above- and belowground characterization is poorly evaluated. In maize, root growth is known to be hindered to a much larger extent than shoot growth and is correlated to photosynthetic activity (Hund et al., 2007). Similar correlations could be expected in soybean and could be a useful tool for large-scale and non-destructive evaluation of the rooting system.

Depending on their target environment, breeders can adopt selection strategy toward specifically adapted cultivars, *i.e.*, cultivars with good performances in a specific set of conditions, or toward broadly adapted cultivars, *i.e.*, cultivars with average or good performances in a large range of conditions (Cooper & Byth, 1996). When broad adaptation is targeted, cultivars are required to be able to express different phenotypes in different environments. This capacity is called phenotypic plasticity. Plastic responses are affected by both inevitable environmental limitations and adaptive adjustments that enhance the organism’s fitness (Sultan, 2000). Interest in selecting directly for adaptive plasticity rather than traits *per se* has risen in the past decade (Gage et al., 2017; Schneider & Lynch, 2020).

Traits can differ in levels of plasticity (Pfennig, 2021). For instance, in sunflower (*Helianthus annuus*), biomass allocation trait plasticity was higher than morphological trait plasticity (Wang et al., 2020). Thus, distinction between performance traits, which contribute directly to fitness (*e.g.*, biomasses), and functional traits, which “impact fitness indirectly via their effects on growth, reproduction and survival” (*e.g.*, root-to-shoot ratio, specific leaf area), should be made (Violle et al., 2007).

In the context of soybean deployment in the north, cold stress at high latitudes is one of the most impactful factor on genotypes’ relative performance in early maturity soybean (Elmerich et al., 2023b). Thus, soybean adaptation to this stress requires unraveling the phenotypic plasticity of traits involved in dark chilling tolerance. In this study, the aims were to (i) highlight plastic traits and their genetic variation in response to dark chilling and to (ii) identify relationships between functional traits involved in dark chilling responses and performance traits to improve breeding strategy.

## 2. Material and Methods

### 2.1. Experimental layout

The experiment was conducted in growth chambers using 80 x 30 x 6 cm rhizotrons (*i.e.*, containers with a transparent wall through which roots can be observed at a regular interval in a non-destructive way). The rhizotrons (©Vienna Scientific Instruments GmbH, Austria) were made of black opaque PVC for the back, sides, and bottom walls and of transparent PVC for the front wall. Drainage holes (0.5 cm in diameter) were made at the bottom. The front wall was covered with black opaque plastic to minimize phototropic response from the roots. The rhizoboxes were inclined at a 40° angle with the front wall down in order to maximize the growth of the roots along the transparent wall.

### 2.2. Plant material and growth conditions

Plants were grown at a 15/9 photoperiod (day/night) at 300 µmol m^-2^ s^-1^ and at a relative humidity of 70% in a mixture of 20% river sand and 80% commercial soil mixture (soil, peat, and compost) (FLOORE, France). The mixture was autoclaved for 20 min at 121°C before filling the rhizotrons.

Two temperature treatments were used: a control treatment and a dark chilling treatment. The control treatment was set at 25/18°C and the dark chilling treatment at 25/10°C (day/night). The transition of temperature between night/day and day/night took an hour, from 7:00 am to 8:00 am and from 10:00 pm to 11:00 pm, respectively. The watering was performed every 2/3 days homogenously at the surface with distilled water in order to maintain the soil moist.

The experiment included five soybean cultivars, two from the “000” maturity group (cv. SIRELIA, cv. ES GOVERNOR) and three from the “00” maturity group (cv. STUMPA, cv. SOPRANA, cv. SULTANA) in six replications per growth chamber. The experimental design was a randomized complete block design (RBBD).

The seeds were inoculated with rhizobium, *Bradyrhizobium japonicum*, (FORCE 48, Lidea Seeds, France) and pregerminated without light in an incubator at 25°C for five days. Seeds with a 0.5 to 1.5 cm radicle were transplanted in the rhizotrons at a depth of 2 cm along the transparent wall. After transplanting the germinated seeds into the rhizoboxes, the two chambers were set at the control treatment temperature. Once the seedlings had reached the emergence stage (VE), one chamber was set at the dark chilling treatment temperature regime. Plants were harvested 29 and 37 days after VE for the dark chilling and the control treatment respectively, once half the plants reached flowering stage (R1). Soybean reaches R1 when one open flower is visible on the main stem at any node (Fehr & Caviness, 1977).

### 2.3. Measurements

#### 2.3.1. Performance traits

At harvest, performance traits attesting for biomass accumulation and allocution were measured at the end of the experiment in the dark chilling condition (25/10°C [day/night]) (**Table 1**). Plant shoots were separated from the roots. The shoots were divided into leaves and stems, and the fresh weights were recorded. The leaf area (LA) was measured using an LI-3100C Area Meter instrument (LI-COR®). The root system was taken out from the soil and carefully washed using demineralized water. The nodules were gently separated from the roots and counted. The nodule-free roots were scanned while being immersed in water using an Epson Perfection V800 scanner. Scan analyses were performed with the WinRhizo Regular V.2016a software (Regent Instruments Inc., Quebec, Canada). Root length and diameter were measured. Each plant compartment was dried at 65°C for 48 hours and weighed.

**Table 1.** Performance traits attesting for biomass accumulation and allocution measured at the end of the experiment in the dark chilling condition (25/10°C [day/night]).

| Traits | Description | Unit |
| --- | --- | --- |
| Biomass (TM) | Total plant dry weight | g |
| Root-to-shoot ratio (R2S) | Root mass divided by shoot mass | no u. |
| Dry matter content (DMC) | Total plant dry weight divided by total plant fresh weight | g g<sup>-1</sup> |
| Leaf area (LA) | Leaf area per plant | cm<sup>2</sup> |
| Plant height (H) | Height of the principal stem from soil surface to meristem | cm |
| Shoot mass (SM) | Shoot dry weight per plant | g |
| Specific leaf area (SLA) | Leaf area divided by leaf dry weight | cm g<sup>-1</sup> |
| Fine root ratio (FRR) | Fine root ($\varnothing < 2\text{mm}$) length divided by total root length | no u. |
| Nodule number (N) | Nodule number per plant | no u. |
| Root diameter (RD) | Average diameter of all roots per plant | cm |
| Root mass (RM) | Root dry weight per plant | mg |
| Total root length (TRL) | Total length of all roots per plant | cm |

Growth rate for the different compartments (shoot, leaf, and root) was calculated as the dry mass divided by the number of days between transplanting and harvest. The elongation rate for leaf and total root was calculated as, respectively, the area and length divided by the number of days between transplanting and harvest. The apparition of new, fully expanded leaf on the main stem was recorded and used to calculate the development rate.

From VE and then once a week, the tap root length was measured. Pictures were taken with a camera (PANASONIC DC-FZ82EF-K) at 90 cm distance from the rhizoboxes and then analyzed with the semi-automated Smart Root (version 4.1) plugin included in the ImageJ (version 1.51) software. The average tap root elongation rate (TRER) was calculated using the following equation (E1). Where, L_t_ and L_t_ are the tap root lengths at day *i+1* and *i*, respectively, t_i+1_ and t_i_ are the days of measurement *i+1* and *i*, respectively, and n is the number of measurements.

The photosynthetic activity parameters were recorded with a MultispeQ device (PHOTOSYNQ INC., USA) on the latest fully developed leaf of the main stem on the middle leaflet. This device measured SPAD and estimated two fluorescence-based photosynthetic parameters using pulse-amplitude modulation (PAM) fluorometry: the maximum quantum efficiency of PSII primary photochemistry (F_v_/F_m_) and the quantum yield electron transport at PSII (ΦPSII) (Maxwell & Johnson, 2000; Kuhlgert et al., 2016). Two measurements were taken at 30 minutes and 7.5 hours after the night period. The first measurement corresponded to t1 and the second to midday. The difference between the measurement at midday and t1 divided by the measurement at midday was calculated. This ratio (R) evaluated the speed of photosynthetic recovery from the night period. All functional traits are described in **Table 2**.

**Table 2.**
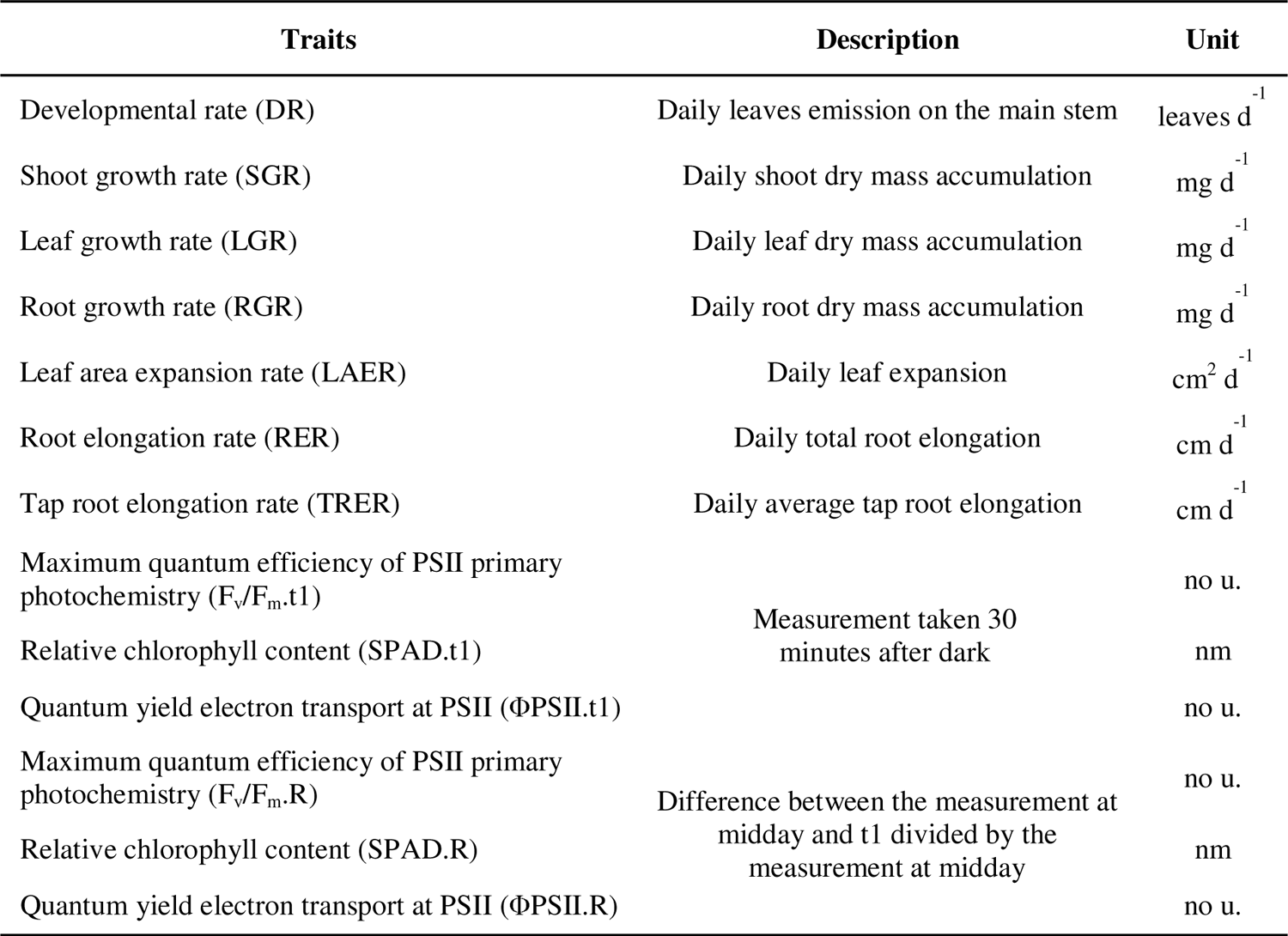
Description of functional traits attesting for growth efficiency measured both in dark chilling (25/10°C [day/night]) and control (25/18°C [day/night]) conditions.

### 2.4. Relative distance plasticity index (RDPI)

The relative distance plasticity index (RDPI) was chosen as a quantitative estimator of the phenotypic plasticity of the functional traits. According to Valladares et al. (2006), this index shows strong statistical power to test for differences in plasticity between genotypes. The RDPI corresponds to the absolute phenotypic distances between individuals of same genotype and different environments, divided by the sum of the two phenotypic values.

Where RDPI*tijk* corresponds to the RDPI of the trait *t* for the cultivar *i* between the repetition *j* from the dark chilling treatment and the repetition *k* from the control treatment, Vdc_tij_ corresponds to the value in the dark chilling treatment of the trait *t* for *j*^th^-repetition of the cultivar *i*, and Vc_tik_ corresponds to the value in the control treatment of the trait *t* for *k*^th^-repetition of the cultivar *i*. RDPI ranges from 0 (no plasticity) to 1 (maximal plasticity).

### 2.5. Statistical analysis

All the analyses were done using Rstudio software (Version 1.2.5033).

The principal component analysis (PCA) was performed using *prcomp* (package: stats) and *fviz_pca_var* (package: factoextra) functions. It included performance and function traits as variables. The *rcorr* function (package: Hmisc) was used to calculate the Pearson’s correlation matrix and the p-value.

For all performance traits measured in the dark chilling treatment, a one-way ANOVA was performed testing the effect of the genotype. If the effect was significant (p-value < 0.05), a multiple comparison test by means of Tukey was done using the *HSD.test* function (package: agricolae). Kruskal–Wallis rank sum test was performed using the *kruskal.test* function (package: stats) followed by Dunn’s test for pairwise multiple comparisons of the ranked data using the *dunn_test* function (package: rstatix). The Bonferonni method was used to adjust p-values for multiple comparisons.

## 3. Results

### 3.1. Performance and functional trait variations and relations under dark chilling conditions

The PCAs of twenty performance and physiological traits of five cultivars under dark chilling conditions explained cumulatively 52.5% of the variation: 35.8% for the first axis and 16.7% for the second (Error! Reference source not found.). Performance traits (**Table 1**) had the highest contributions and were correlated with the first axis while functional traits were discriminated by the second axis. Above- and belowground performance traits were mainly positively correlated except for the specific leaf area and the fine root ratio. For example, the leaf area and the shoot biomass showed positive correlation with the average root diameter (r = 0.45 and r = 0.50, respectively). The fine root ratio showed negative correlation with other performance traits such as the total biomass (r = −0.41). The SPAD measured after the dark period (t1) was negatively correlated with the specific leaf area (r = −0.59). The recuperation capacity (R) of the quantum yield electron transport at PSII (ΦPSII [R]) was negatively correlated with aboveground performance traits (r = −0.42, r = −0.44, r = −0.63, for shoot mass, leaf mass, and leaf area, respectively). Photosynthetic activity traits (t1) were positively correlated with plant height (r = 0.44, r = 0.50, for ΦPSII and SPAD, respectively) and the TRER (r = 0.67 for SPAD).

### 3.2. Genetic variation of performance traits in responses to dark chilling

Shoot biomass and leaf area are two major performance traits in terms of production and accumulation of organic matter showing a positive correlation with yield and yield components. Significant differences were observed between the genotypes for these two performance traits in the dark chilling conditions (p-value < 0.05). Cv. SOPRANA showed the highest shoot biomass while cv. SIRELIA had the lowest (Figure 2a). Cv. RGT STUMPA had the greatest leaf area whereas cv. SIRELIA had the lowest (Figure 2b). Genotypic differences were observed for other performance traits (**Table 3**).

**Figure 1.**
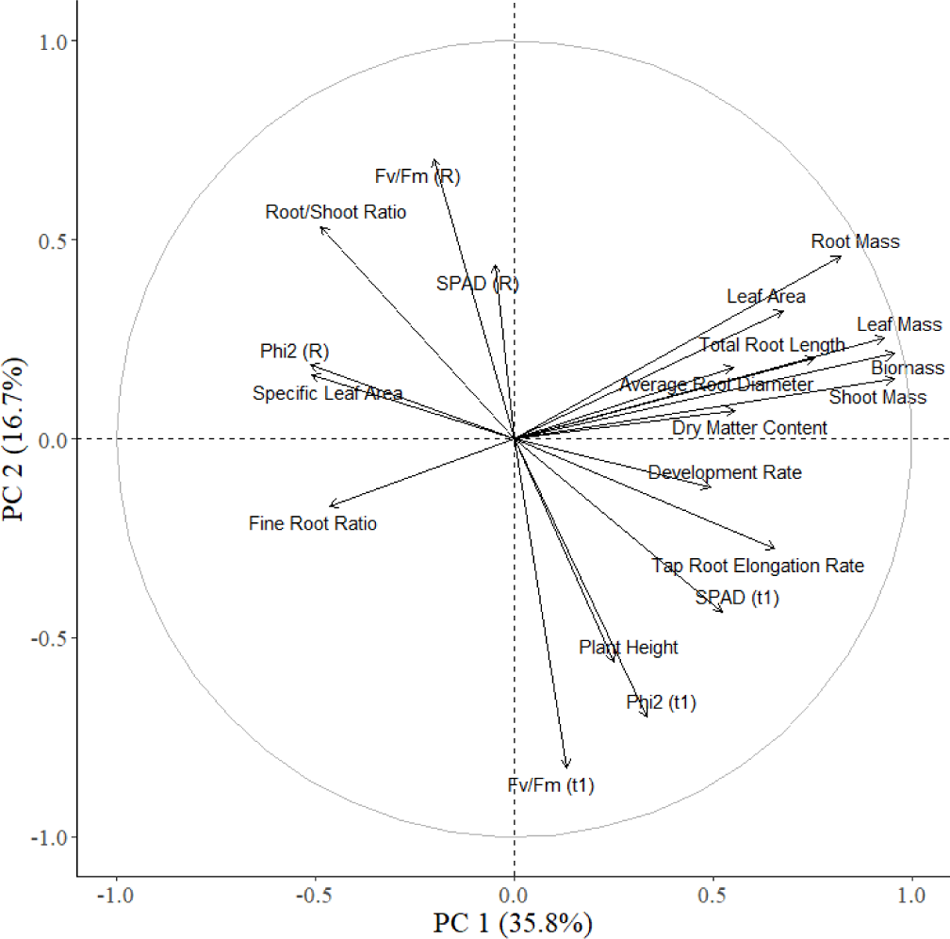
Principal component analyses of twenty functional and performance traits under dark chilling condition (25/10°C [day/night]). The two principal components (PC 1 and PC 2) explained 52.5% of the variation.

**Figure 2.**
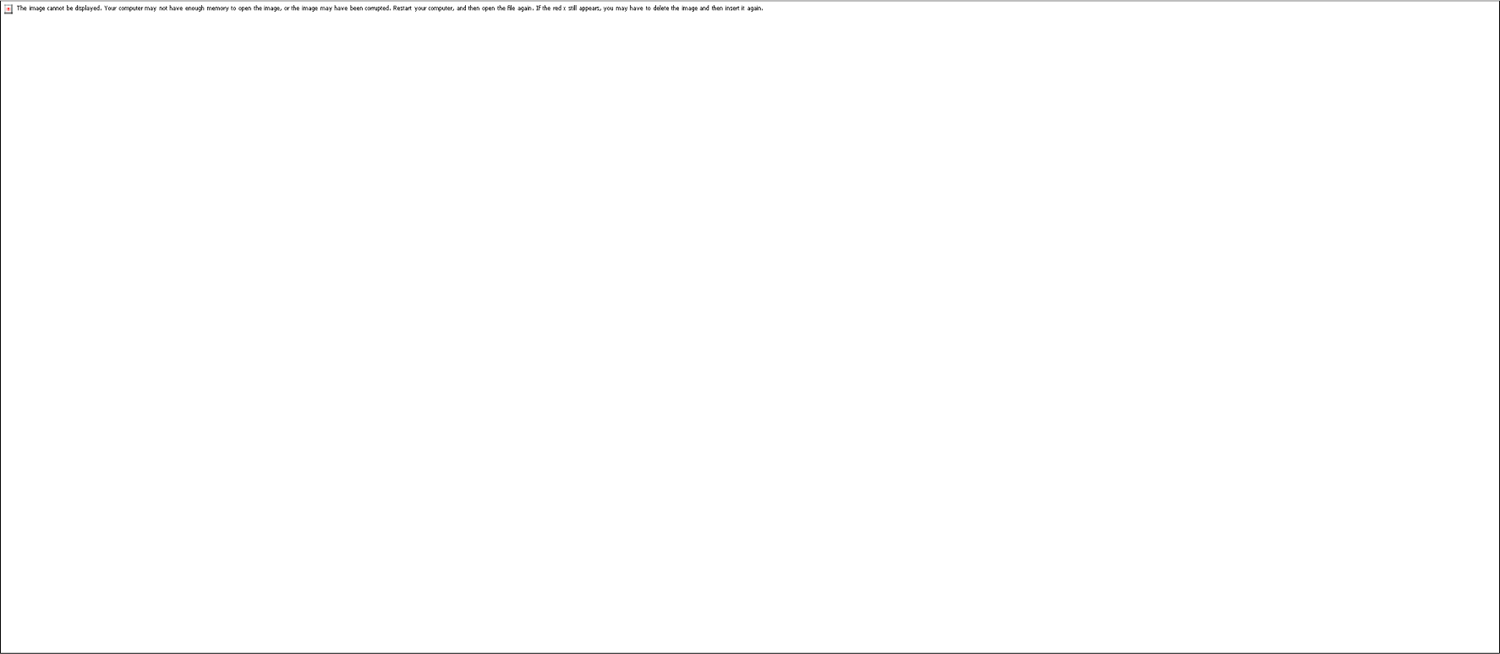
Performance traits (**(a)** shoot mass, **(b)** leaf area) of five cultivars (cv. ES GOVERNOR, cv. RGT STUMPA, cv. SIRELIA, cv. SOPRANA, and cv. SULTANA) under dark chilling stress (25/10°C [day/night]). The letter above each error bar (standard error) corresponds to the result of the HSD test. Genotypes with a common letter were not significantly different at the 5% level of significance.

**Table 3.** Means and standard deviations of the twelve performance traits for five genotypes measured in dark chilling stress (25/10°C [day/night]). For each trait, genotype means followed by a common letter are not significantly different, according to the HSD test at the 5% level of significance.

| Traits | *p*-value | ES GOVERNOR | RGT STUMPA | SIRELIA | SOPRANA | SULTANA |
| --- | --- | --- | --- | --- | --- | --- |
| Biomass (g) | 0.019* | 12.9 ± 2.8<sup>ab</sup> | 12.2 ± 2.1<sup>ab</sup> | 10.2 ± 2.0<sup>b</sup> | 15.4 ± 0.9<sup>a</sup> | 11.7 ± 1.1<sup>ab</sup> |
| Root-to-Shoot | 0.013* | 0.26 ± 0.05<sup>abc</sup> | 0.32 ± 0.06<sup>a</sup> | 0.31 ± 0.06<sup>ab</sup> | 0.25 ± 0.02<sup>c</sup> | 0.25 ± 0.04<sup>bc</sup> |
| Dry matter content (g/g) | 0.000*** | 0.22 ± 0.02<sup>a</sup> | 0.19 ± 0.01<sup>b</sup> | 0.16 ± 0.01<sup>c</sup> | 0.20 ± 0.01<sup>ab</sup> | 0.20 ± 0.01<sup>ab</sup> |
| Leaf area (cm<sup>2</sup>) | 0.039* | 1 657 ± 247<sup>ab</sup> | 2 093 ± 230<sup>a</sup> | 1 301 ± 129<sup>b</sup> | 1 757 ± 227<sup>ab</sup> | 1 804 ± 416<sup>ab</sup> |
| Plant height (cm) | 0.009** | 38.8 ± 1<sup>b</sup> | 45.6 ± 2.2<sup>b</sup> | 40.8 ± 6.2<sup>b</sup> | 97.5 ± 10.8<sup>a</sup> | 38.5 ± 5.4<sup>b</sup> |
| Shoot mass (g) | 0.043* | 10.1 ± 2.6<sup>ab</sup> | 9.07 ± 0.9<sup>ab</sup> | 7.9 ± 2.3<sup>b</sup> | 12.4 ± 2.1<sup>a</sup> | 9.3 ± 0.8<sup>ab</sup> |
| Specific leaf area (cm<sup>2</sup>/g) | 0.013* | 264 ± 36<sup>b</sup> | 291 ± 61<sup>b</sup> | 298 ± 38<sup>ab</sup> | 261 ± 14<sup>b</sup> | 385 ± 72<sup>ab</sup> |
| Fine Root ratio | 0.008** | 0.9990 ± 5.10<sup>-4a</sup> | 0.9973 ± 5.10<sup>-4b</sup> | 0.9991 ± 3.10<sup>-4a</sup> | 0.9986 ± 5.10<sup>-4ab</sup> | 0.9982 ± 1.10<sup>-4ab</sup> |
| Nodule number | 0.013* | 12 ± 7.26<sup>ab</sup> | 11.8 ± 7.82<sup>ab</sup> | 9.5 ± 4.5<sup>b</sup> | 28 ± 6<sup>ab</sup> | 32 ± 12.17<sup>a</sup> |
| Root diameter (cm) | 0.016* | 2.33 ± 0.18<sup>ab</sup> | 2.61 ± 0.2<sup>a</sup> | 2.05 ± 0.17<sup>b</sup> | 2.42 ± 0.05<sup>ab</sup> | 2.31 ± 0.38<sup>ab</sup> |
| Root mass (g) | 0.063ns | 2.72 ± 0.98 | 3.26 ± 0.52 | 2.33 ± 0.44 | 2.98 ± 0.41 | 2.39 ± 0.57 |
| Total root length (cm) | 0.045* | 17 752 ± 3 140<sup>ab</sup> | 15 493 ± 2 370<sup>ab</sup> | 14 609 ± 3 260<sup>ab</sup> | 19 666 ± 3 793<sup>a</sup> | 12 213 ± 1 857<sup>b</sup> |
| Fv/Fm.t1 | 0.706ns | 0.68 ± 0.02 | 0.68 ± 0.03 | 0.69 ± 0.02 | 0.70 ± 0.01 | 0.69 ± 0.03 |
| ΦPSII.t1 | 0.545ns | 0.52 ± 0.05 | 0.50 ± 0.02 | 0.50 ± 0.06 | 0.54 ± 0.03 | 0.51 ± 0.04 |
| SPAD.t1 (nm) | 0.068ns | 43.1 ± 2.83 | 44.7 ± 4.82 | 39.3 ± 4.87 | 46.2 ± 2.23 | 41.0 ± 1.73 |
| Fv/Fm.R | 0.679ns | 0.05 ± 0.03 | 0.04 ± 0.04 | 0.05 ± 0.02 | 0.02 ± 0.01 | 0.04 ± 0.03 |
| ΦPSII.R | 0.480ns | 0.06 ± 0.05 | 0.05 ± 0.04 | 0.09 ± 0.05 | 0.07 ± 0.02 | 0.04 ± 0.03 |
| SPAD.R (nm) | 0.630ns | 0.04 ± 0.02 | 0.02 ± 0.02 | 0.04 ± 0.04 | 0.04 ± 0.03 | 0.03 ± 0.03 |

### 3.3. Phenotypic plasticity of traits in response to dark chilling

The average RDPI of the 16 functional traits ranged from almost 0 to 0.48 (**Table 4**). SPAD, F_v_/F_m_ and ΦPSII showed the highest RDPI when considering the recuperation capacity (R) but the lowest for their measurement after dark (t1). Growth rates of the different compartments had similar RDPIs, around 0.18.

**Table 4.** Means and standard deviations of the relative distance plasticity index (RDPI) of functional traits for five genotypes. The traits are displayed by mean RDPI. Level of significance of the genotypic effect was tested by a Kruskal–Wallis test and the p-value was adjusted by the Bonferroni method. For each trait, genotype means followed by a common letter were not significantly different according to the Dunn test at the 5% level of significance.

| Traits | Mean RDPI | *p*-value | ES GOVERNOR | RGT STUMPA | SIRELIA | SOPRANA | SULTANA |
| --- | --- | --- | --- | --- | --- | --- | --- |
| SPAD.R | 0.48 ± 0.28 | < 0.001*** | 0.34 ± 0.20b | 0.65 ± 0.24a | 0.51 ± 0.33ab | 0.37 ± 0.19b | 0.50 ± 0.30a |
| Fv/Fm.R | 0.48 ± 0.26 | < 0.001*** | 0.48 ± 0.25ab | 0.60 ± 0.27a | 0.48 ± 0.24ab | 0.37 ± 0.22b | 0.45 ± 0.29a |
| ΦPSII.R | 0.39 ± 0.23 | 0.133 ns | 0.37 ± 0.21 | 0.34 ± 0.22 | 0.47 ± 0.23 | 0.31 ± 0.22 | 0.39 ± 0.25 |
| Root Growth Rate | 0.19 ± 0.17 | 0.002** | 0.3 ± 0.23a | 0.20 ± 0.14ab | 0.11 ± 0.08b | 0.28 ± 0.18a | 0.12 ± 0.08ab |
| Shoot Growth Rate | 0.18 ± 0.16 | < 0.001*** | 0.28 ± 0.20ab | 0.17 ± 0.12ab | 0.15 ± 0.14bc | 0.28 ± 0.15a | 0.06 ± 0.06c |
| Leaf Growth Rate | 0.18 ± 0.15 | < 0.001*** | 0.26 ± 0.20a | 0.16 ± 0.10a | 0.15 ± 0.14ab | 0.27 ± 0.15a | 0.06 ± 0.06b |
| Root Elongation Rate | 0.18 ± 0.14 | 0.002** | 0.24 ± 0.18ab | 0.14 ± 0.09ab | 0.12 ± 0.07b | 0.29 ± 0.19a | 0.12 ± 0.08b |
| Leaf Area Elongation Rate | 0.17 ± 0.12 | 0.317 ns | 0.20 ± 0.14 | 0.19 ± 0.14 | 0.18 ± 0.11 | 0.16 ± 0.09 | 0.13 ± 0.08 |
| Specific Leaf Area | 0.13 ± 0.10 | 0.028* | 0.17 ± 0.08a | 0.14 ± 0.08ab | 0.11 ± 0.06ab | 0.17 ± 0.18ab | 0.09 ± 0.08b |
| Dry Matter Content | 0.13 ± 0.06 | < 0.001*** | 0.18 ± 0.05a | 0.10 ± 0.05b | 0.1 ± 0.05b | 0.13 ± 0.06b | 0.13 ± 0.03ab |
| Tap Root Elongation Rate | 0.12 ± 0.09 | < 0.001*** | 0.13 ± 0.06a | 0.05 ± 0.04b | 0.10 ± 0.10ab | 0.17 ± 0.10a | 0.16 ± 0.08a |
| Root to Shoot Ratio | 0.10 ± 0.07 | 0.06 ns | 0.10 ± 0.07 | 0.09 ± 0.06 | 0.13 ± 0.08 | 0.07 ± 0.06 | 0.08 ± 0.05 |
| Development Rate | 0.09 ± 0.05 | 0.051 ns | 0.09 ± 0.05 | 0.08 ± 0.04 | 0.10 ± 0.05 | 0.07 ± 0.05 | 0.11 ± 0.06 |
| SPAD.t1 | 0.06 ± 0.04 | < 0.001*** | 0.06 ± 0.03ab | 0.09 ± 0.04a | 0.06 ± 0.04ab | 0.05 ± 0.03b | 0.04 ± 0.04b |
| ΦPSII.t1 | 0.06 ± 0.03 | 0.017* | 0.06 ± 0.04ab | 0.07 ± 0.03a | 0.05 ± 0.03ab | 0.04 ± 0.03b | 0.06 ± 0.04ab |
| Fv/Fm.t1 | 0.03 ± 0.02 | < 0.001*** | 0.05 ± 0.02a | 0.04 ± 0.02ab | 0.03 ± 0.01bc | 0.01 ± 0.01c | 0.03 ± 0.02bc |

For the majority of the considered traits, the genotypic effect was significant (**Table 4**). Cv. RGT STUMPA showed significantly higher RDPI for SPAD and F_v_/F_m_ ratios than cv. SOPRANA (Figure 3a **and 3b**). Interestingly, cv. SOPRANA had the highest RDPI for shoot growth rate (Figure 3c). Cv. RGT STUMPA presented low RDPI for TRER (Figure 3d).

**Figure 3.**
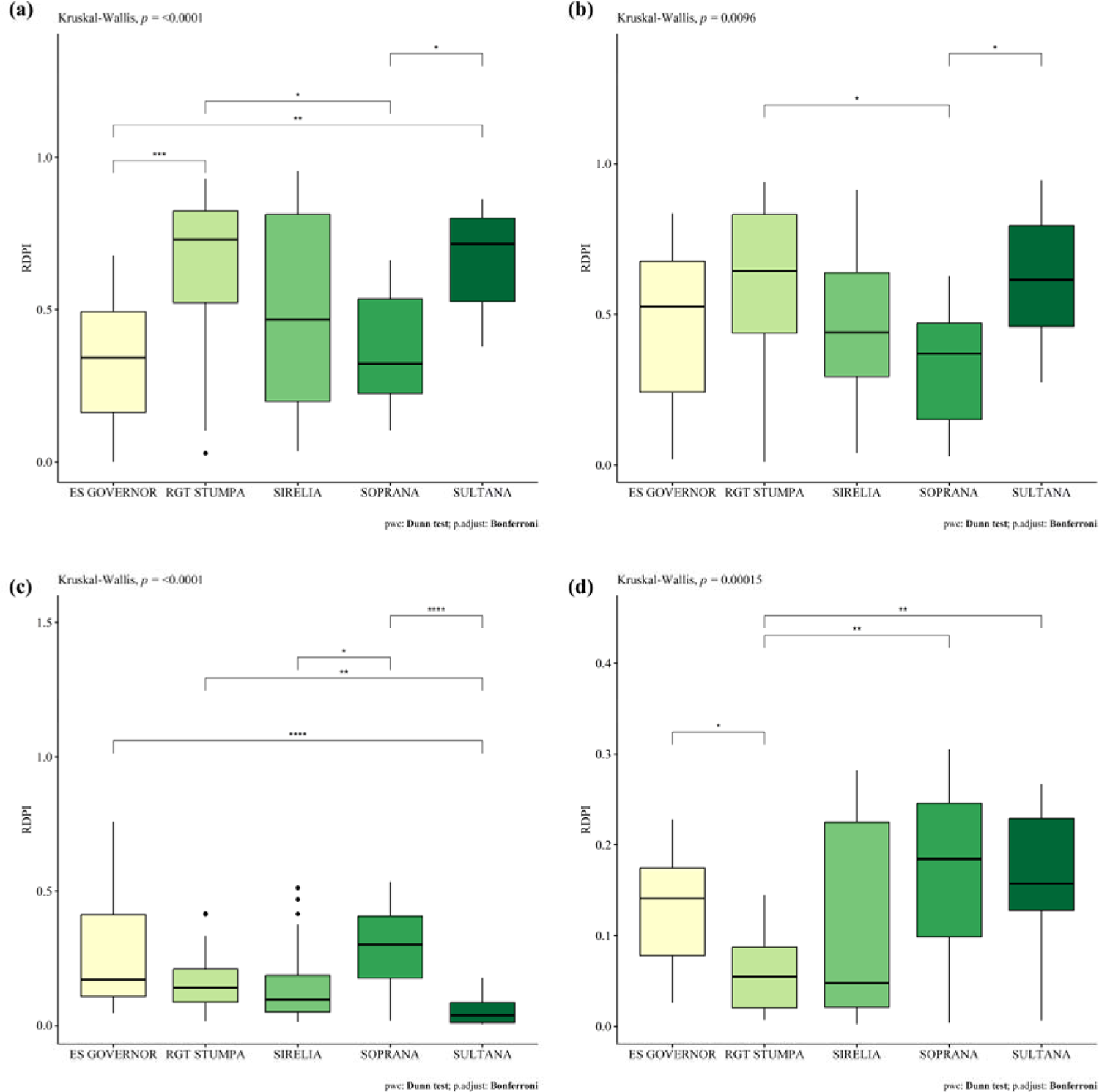
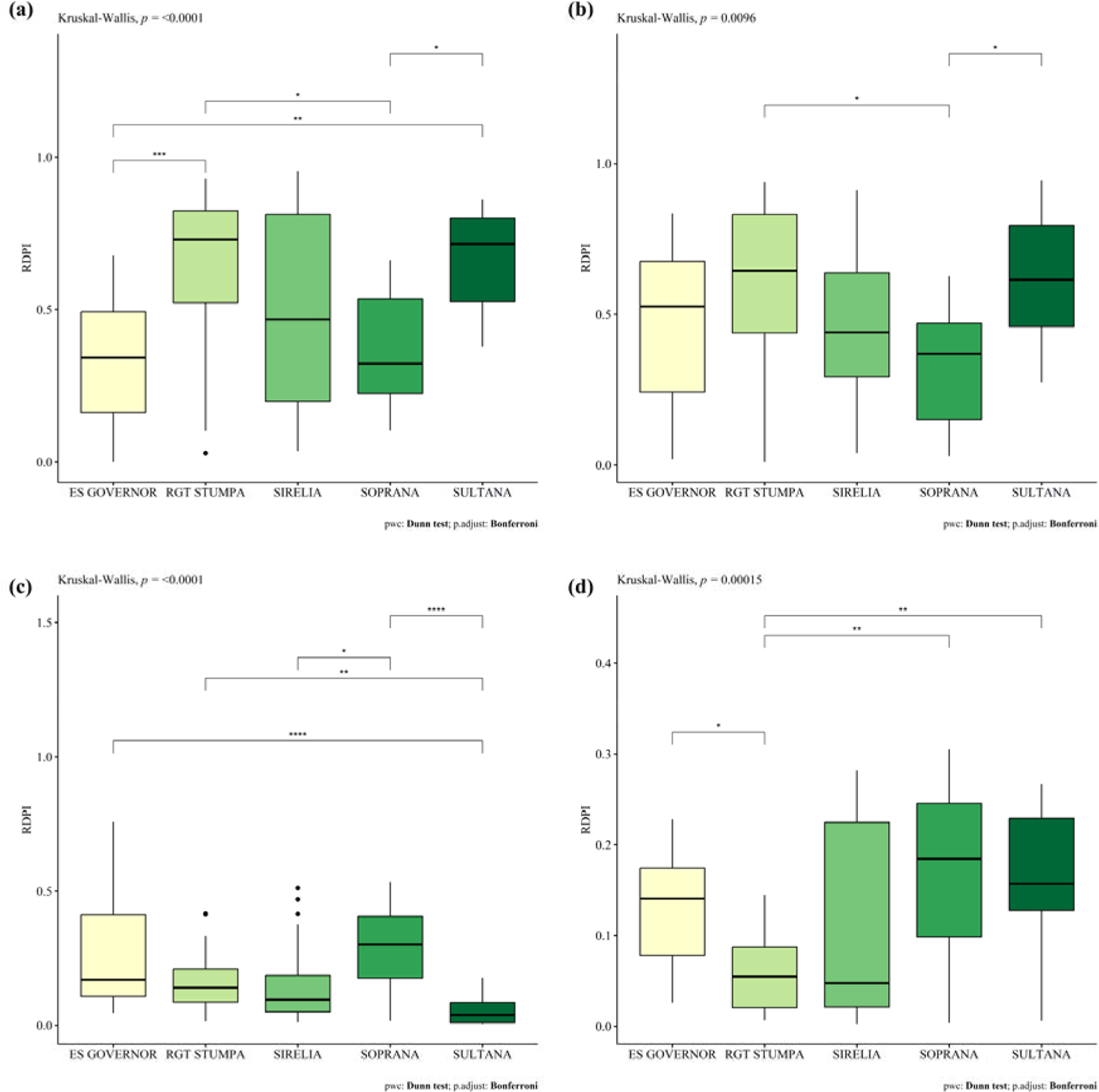
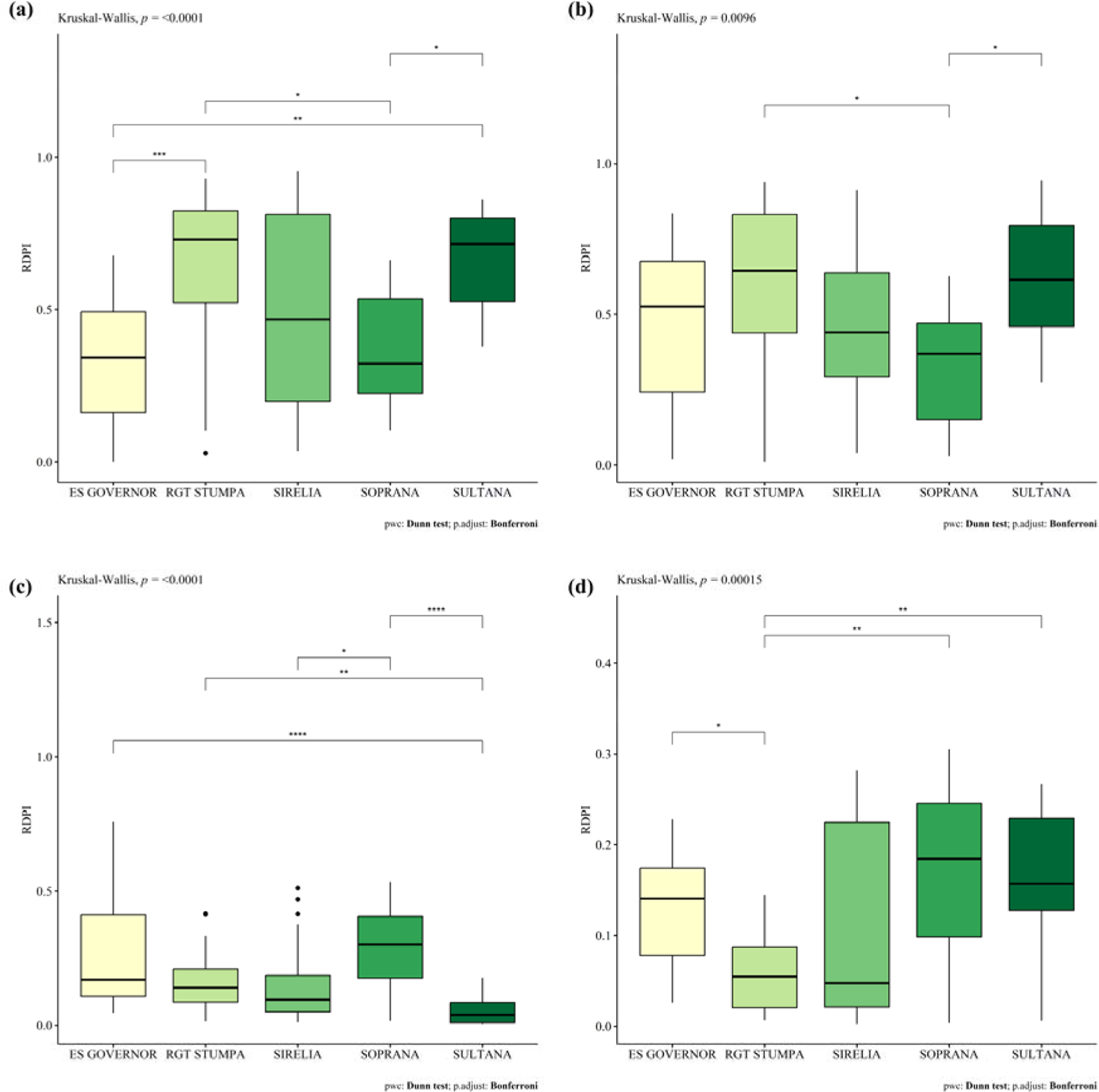
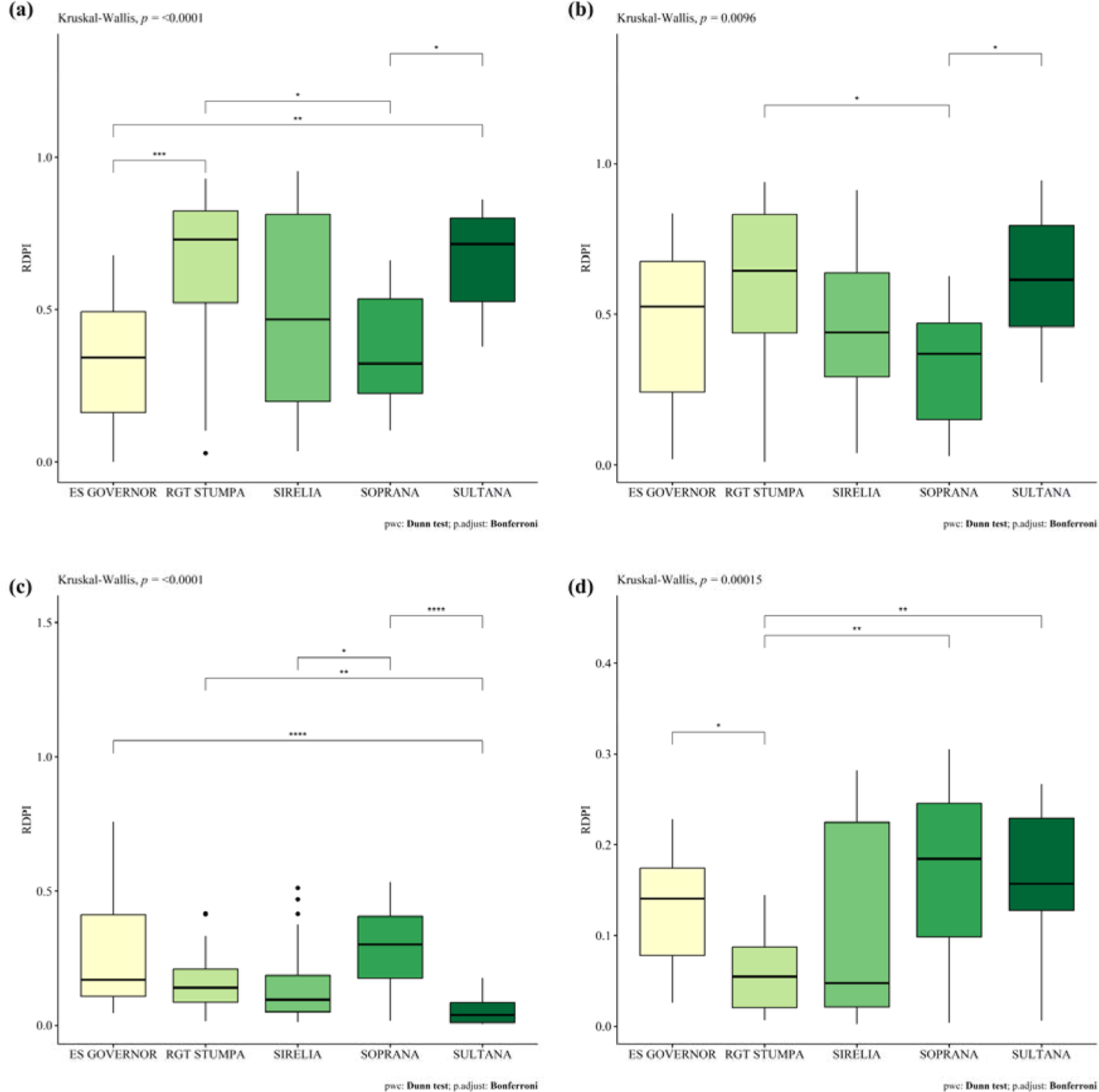
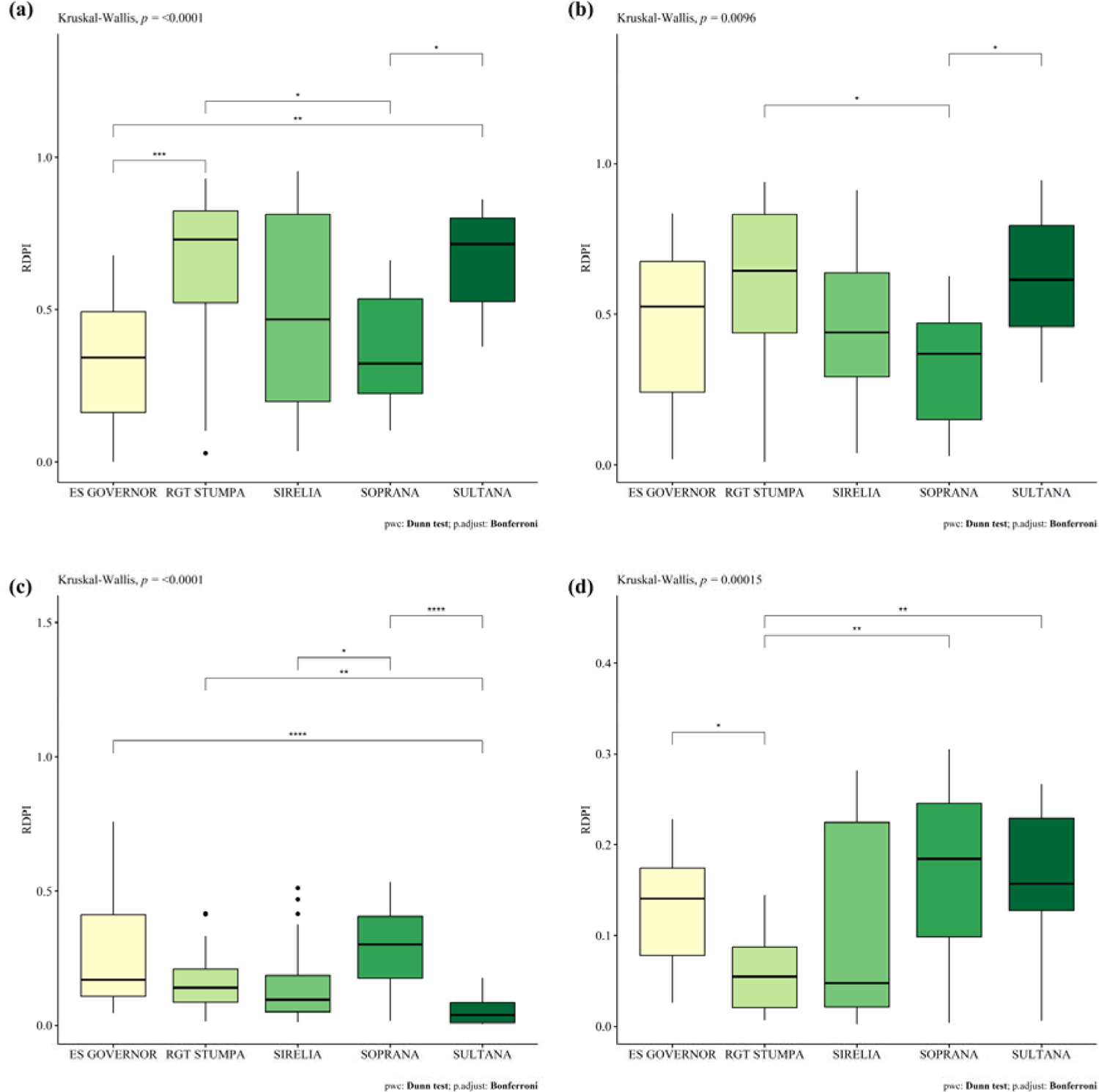
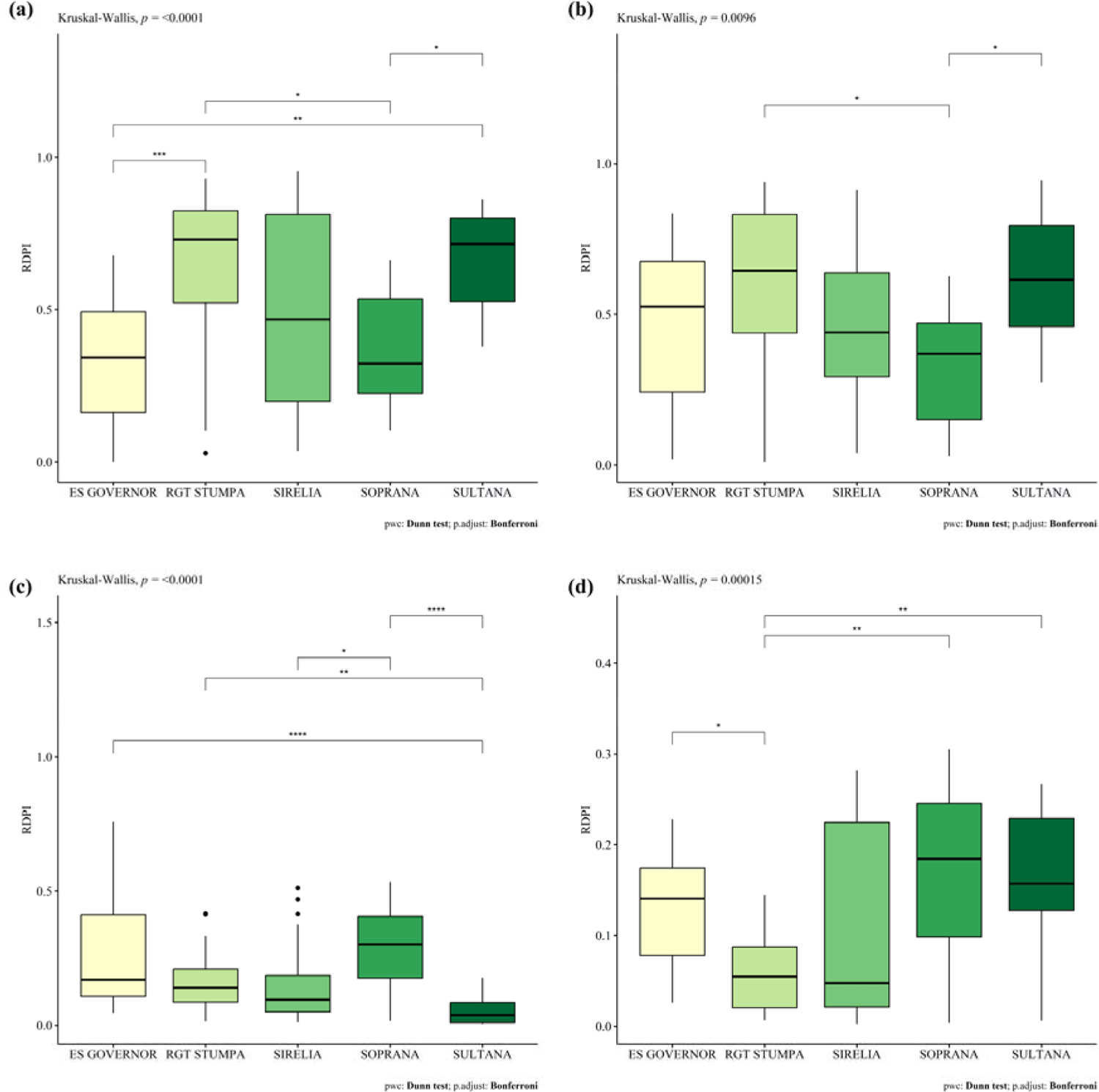
Relative distance plasticity index (RDPI) for (a) SPAD.R, (b) F_v_/F_m_.R, (c) shoot growth rate, and (d) tap root elongation rate. Kruskal–Wallis test statistics are displayed on the top left and the pairwise comparison test is indicated in the bottom right.

## 4. Discussion

### 4.1. Plasticity of soybean physiological, morphological, and performance traits to dark chilling

Phenotypic plasticity plays a key role in plants’ capacity to adapt to varying environments and in their chances of overcoming environmental constraints, such as dark chilling stress (Merilä & Hendry, 2014). Herein, from a panel of modern early soybean cultivars, different levels of plasticity were observed between functional traits. These results suggested that even though plasticity was not a parameter intentionally targeted by breeders, it was kept within the modern germplasm. According to Brooker et al. (2022), this assessment of existing plasticity is essential if the phenotypic plasticity of a trait is desired to become a breeding target. In soybean, photosynthetic activity recovery traits showed on average the highest plasticity indexes (*ca.* 0.45 ± 0.04), followed by biomass production traits (*ca.* 0.18 ± 0.005) and morphological traits (*ca*. 0.11 ± 0.02). Photosynthetic activity after dark had the lowest plasticity indices (*ca.* 0.05 ± 0.01).

Shoot and root biomass showed highly positive correlation but low to no differences between cultivars, respectively. Thus, an allometric equation could be used to predict root biomasses that remain challenging to measure in field experiments (Zhu et al., 2011). However, shoot and root growth rate showed an average level of plasticity and highly significant differences between cultivars. These could be used to discriminate between cultivars and explore their tolerance to dark chilling stress (Rivera et al., 2021).

The root-to-shoot ratio is often considered to reflect the crop response to abiotic influences and to be a marker of the trade-off between above- and belowground biomass allocation (Poorter & Sack, 2012). Thus, it is generally used to predict the root biomass more accurately than when an equation derived from the root and shoot biomass is used (Mokany et al., 2006). Surprisingly, under dark chilling stress, an increase in soybean root-to-shoot ratio resulted in a smaller shoot biomass, but the root biomass was independent. Although differences between cultivars in root-to-shoot ratio were observed, this morphological trait had a low plasticity index that was similar among cultivars. One interpretation could be that soybean cultivars under dark chilling stress had a similar strategy of greater partitioning to the root system due to lower demand for photosynthates from aboveground parts experiencing lower growth rates (Reddy et al., 2017; Walne & Reddy, 2022). The preferred allocation of photosynthates to root biomass might confer greater resilience to the genotypes after a stressed period.

Physiological processes are known to be highly sensitive to environmental cues (Allen & Ort, 2001). Photosynthetic traits measured after the night period were positively correlated to each other while no correlation was observed between photosynthesis recovery traits. This result suggested that the photosynthesis recovery dynamics depend on the considered trait and its related function. Further investigations are needed to assess the pattern of photosynthesis after dark in soybean. In the literature, instantaneous or delayed photosynthesis dysfunction have been observed in crops of tropical and subtropical origin such as tomato, *Lycopersicon esculentum,* (Martin et al., 1981), mango, *Mangifera indica* L., (Allen et al., 2000) or maize, *Zea mays* L., (Long et al., 1983).

In our study, the TRER was positively correlated to the biomass production but also to the photosynthetic activity after the dark period. Thus, higher photosynthesis after dark led to a higher elongation rate, suggesting that photosynthates are used initially for root cell expansion. In optimum conditions, roots elongate at night, when the entire plant is decreasing in dry mass, because of carbon use in respiration (Lambers & Oliveira, 2019). Thus, this could be validated through CO_2_ labelling to follow carbon flux in the plant after the dark period (Epron et al., 2012). Finally, our results allow for the proposition that photosynthetic activity could be used to assess the dynamic of soybean tap root under dark chilling stress.

Among the identified plastic traits, physiological traits such as the photosynthetic activity recovery (F_v_/F_m_.R and SPAD.R) and growth traits such as the shoot growth rate and the TRER appeared suitable to study the difference of plastic response to dark chilling stress between cultivars. Outdoor experiments are required to assess if these results are transposable to the field (Zhu et al., 2011).

### 4.2. The phenotypic plasticity of traits involved in dark chilling response and their genetic variation

Chlorophyll fluorescence parameters were reported to allow for discrimination of soybean cultivars under dark chilling stress (Van Heerden et al., 2003; Strauss et al., 2006; Krüger et al., 2014), but no genotypic difference was observed for photosynthetic traits in our experiment. One explanation could be that, contrary to these studies, only temperate soybean cultivars from early maturity groups were used. A solution could be to include varied maturity groups and cultivar origins to confront the use of chlorophyll fluorescence traits or their plasticity to discriminate between genotypes.

Under dark chilling stress, cv. SOPRANA and cv. SULTANA showed high performance in shoot growth and shoot performance (high shoot mass and leaf area) (Figure 2) but they had contrasting phenotypic plasticity, suggesting that two behaviors could lead to the same biomass production at flowering. On one hand, cv. SULTANA growth performance was associated with a large phenotypic plasticity in physiological traits (*i.e*., F_v_/F_m_.R and SPAD.R) but a small one in aerial performance traits (*i.e.*, shoot growth rate) (Figure 3a**, 3b, 3c**). On the other hand, cv. SOPRANA growth performance was associated with a small phenotypic plasticity in physiological traits but a large one in aerial performance traits (Figure 3a**, 3b, 3c**). For the two cultivars, high phenotypic plasticity in TRER was observed. Cv. SOPRANA behavior might indicate a greater stability and, likely, a greater tolerance to dark chilling stress. Further experiments should focus on assessing active adaptive plasticity, corresponding to plasticity that increases crop fitness or survival, to confirm this hypothesis (Acasuso-Rivero et al., 2019).

When screening for large panels of genotypes and the implementation of a trait measurement in a breeding program, attention should be given to the feasibility and cost of the measurements (Reynolds et al., 2019). This is even more true when selecting for phenotypic plasticity, as it requires the measurement of the trait in at least two environments, in our case with contrasted temperature regimes (Lobet et al., 2019). In our study, the photosynthetic activity recovery showed both high plasticity and significant difference between genotypes. Moreover, it was correlated with the TRER, which could also be a targeted trait by breeder (Schneider & Lynch, 2020). The methodology used in this study to measure the photosynthetic activity presented several advantages. It was fast (20 seconds per measurements), easy, non-destructive, and cheap compared to other investments used in plant breeding (Heffner et al., 2010). Thus, we recommend the use of photosynthetic activity recovery trait plasticity as a proxy for dark chilling tolerance in soybean.

Finally, although genotypes from the same crop may share patterns of plasticity for certain traits, they may also differ in the amount, direction, and timing of plastic responses to a given environmental cue, such as dark chilling (Sultan, 2000). Our results suggested an existing genetic differentiation between cultivars leading to different pattern of plastic response. The plastic traits identified in this study (*i.e.,* the recovery of photosynthetic activity, the growth rates, and the TRER) should be validated *in situ* and could be used to screen a larger panel of cultivars for their tolerance to dark chilling and to enhance soybean adaptation to northern latitudes.

## 5. Conclusion

Dark chilling stress is a major limitation for European soybean expansion northward. Thus, it requires both adaptive and resilient cultivars to weather these conditions. Our results indicated a strong positive correlation between above- and belowground biomass during soybean vegetative growth under dark chilling. It suggests that, for breeding purpose like genotype screening, the evaluation of aboveground biomass was sufficient and informative of the belowground traits under dark chilling stress. As belowground traits remained difficult to measure in large-scale evaluation as observed in breeding programs, these results could guide breeders toward the evaluation of shoot biomass as a proxy for crop biomass accumulation during soybean vegetative growth under dark chilling stress. Moreover, chlorophyll fluorescence traits commonly used as a proxy for cultivars’ tolerance to abiotic stresses did not discriminate early soybean cultivars in our experimental conditions, but their plasticity did. Among performance, morphological, and physiological traits, different levels of plasticity were observed. The plasticity of the shoot growth rate, the TRER, and the photosynthetic activity recovery traits were interesting for studying the response of soybean genotypes to dark chilling stress. Two contrasted trait plasticity patterns were used by the cultivars to achieve high biomass production. Finally, perspectives of this work would be to assess the role of the identified plastic traits in both crop fitness and yield under field conditions. The plastic traits could also be used to screen a larger panel of genotypes for their tolerance to dark chilling stress.

## Acknowledgements

The authors would like to thanks Céline Roisin, Aurore Couturier, Issifou Amadou and Alban Muller for their assistance in the experiment. They acknowledge AGHYLE laboratory for providing the rhizotrons and the controlled chambers for the experiment.

## Authors contribution

CE, GB, BL and MPF designed the experiment. CE, GB and BL conducted the experiment and collected data. CE analysed the data. CE, BL and MPF wrote the first manuscript draft. All author participated to the revision of the manuscript.

